# Fine-mapping of an expanded set of type 2 diabetes loci to single-variant resolution using high-density imputation and islet-specific epigenome maps

**DOI:** 10.1101/245506

**Authors:** Anubha Mahajan, Daniel Taliun, Matthias Thurner, Neil R Robertson, Jason M Torres, N William Rayner, Valgerdur Steinthorsdottir, Robert A Scott, Niels Grarup, James P Cook, Ellen M Schmidt, Matthias Wuttke, Chloé Sarnowski, Reedik Mägi, Jana Nano, Christian Gieger, Stella Trompet, Cécile Lecoeur, Michael Preuss, Bram Peter Prins, Xiuqing Guo, Lawrence F Bielak, DIAMANTE Consortium, Amanda J Bennett, Jette Bork-Jensen, Chad M Brummett, Mickaël Canouil, Kai-Uwe Eckardt, Krista Fischer, Sharon LR Kardia, Florian Kronenberg, Kristi Läll, Ching-Ti Liu, Adam E Locke, Jian′an Luan, Ioanna Ntalla, Vibe Nylander, Sebastian Sch࿆nherr, Claudia Schurmann, Loïc Yengo, Erwin P Bottinger, Ivan Brandslund, Cramer Christensen, George Dedoussis, Jose C Florez, Ian Ford, Oscar H Franco, Timothy M Frayling, Vilmantas Giedraitis, Sophie Hackinger, Andrew T Hattersley, Christian Herder, M Arfan Ikram, Martin Ingelsson, Marit E Jørgensen, Torben Jørgensen, Jennifer Kriebel, Johanna Kuusisto, Symen Ligthart, Cecilia M Lindgren, Allan Linneberg, Valeriya Lyssenko, Vasiliki Mamakou, Thomas Meitinger, Karen L Mohlke, Andrew D Morris, Girish Nadkarni, James S Pankow, Annette Peters, Naveed Sattar, Alena Stančáková, Konstantin Strauch, Kent D Taylor, Barbara Thorand, Gudmar Thorleifsson, Unnur Thorsteinsdottir, Jaakko Tuomilehto, Daniel R Witte, Josée Dupuis, Patricia A Peyser, Eleftheria Zeggini, Ruth J F Loos, Philippe Froguel, Erik Ingelsson, Lars Lind, Leif Groop, Markku Laakso, Francis S Collins, J Wouter Jukema, Colin N A Palmer, Harald Grallert, Andres Metspalu, Abbas Dehghan, Anna Köttgen, Goncalo Abecasis, James B Meigs, Jerome I Rotter, Jonathan Marchini, Oluf Pedersen, Torben Hansen, Claudia Langenberg, Nicholas J Wareham, Kari Stefansson, Anna L Gloyn, Andrew P Morris, Michael Boehnke, Mark I McCarthy

## Abstract

We aggregated genome-wide genotyping data from 32 European-descent GWAS (74,124 T2D cases, 824,006 controls) imputed to high-density reference panels of >30,000 sequenced haplotypes. Analysis of ˜27M variants (˜21M with minor allele frequency [MAF]<5%), identified 243 genome-wide significant loci (*p*<5×10^−8^; MAF 0.02%-50%; odds ratio [OR] 1.04-8.05), 135 not previously-implicated in T2D-predisposition. Conditional analyses revealed 160 additional distinct association signals (*p*<10^−5^) within the identified loci. The combined set of 403 T2D-risk signals includes 56 low-frequency (0.5%≤MAF<5%) and 24 rare (MAF<0.5%) index SNPs at 60 loci, including 14 with estimated allelic OR>2. Forty-one of the signals displayed effect-size heterogeneity between BMI-unadjusted and adjusted analyses. Increased sample size and improved imputation led to substantially more precise localisation of causal variants than previously attained: at 51 signals, the lead variant after fine-mapping accounted for >80% posterior probability of association (PPA) and at 18 of these, PPA exceeded 99%. Integration with islet regulatory annotations enriched for T2D association further reduced median credible set size (from 42 variants to 32) and extended the number of index variants with PPA>80% to 73. Although most signals mapped to regulatory sequence, we identified 18 genes as human validated therapeutic targets through coding variants that are causal for disease. Genome wide chip heritability accounted for 18% of T2D-risk, and individuals in the 2.5% extremes of a polygenic risk score generated from the GWAS data differed >9-fold in risk. Our observations highlight how increases in sample size and variant diversity deliver enhanced discovery and single-variant resolution of causal T2D-risk alleles, and the consequent impact on mechanistic insights and clinical translation.

---

The analysis of array-based genome-wide association studies (GWAS) has, for the past decade, provided the most powerful approach to identify genetic variants contributing to risk of complex traits such as type 2 diabetes (T2D). This approach has provided robust detection of many thousands of associated regions across many hundreds of traits, including >120 loci influencing risk of T2D ^1-3^. Conversion of these genetic discoveries into mechanistic and translational insights has often been challenged by the incomplete coverage of the arrays used for primary genotyping, imperfect performance of the reference panels available for imputation, extensive local linkage disequilibrium (LD), and inadequate sample sizes. This has implications for power to detect low-frequency alleles with population-scale impact, to deliver clinically-relevant risk prediction, and to define molecular mechanisms involved in disease predisposition. In the present study, we address limitations of previous studies by combining GWAS data from ˜900,000 European participants with dense, high-quality imputation to provide the most comprehensive view to date of the genetic contribution to T2D with respect to locus discovery, causal variant resolution, and mechanistic insight.

## RESULTS

### Study overview

We combined genome-wide association data from 32 studies, including 74,124 T2D cases and 824,006 controls of European ancestry (effective sample size 231,436). This represents a 3.2-fold increase in effective sample size from the previous largest genome-wide study of T2D risk in Europeans^1^. With this sample size, assuming accurate imputation (imputation quality score >0.8), we had >80% power to detect T2D association (at α=5×10^−8^) with variants of minor allele frequency (MAF) ≥5% and odds ratio (OR) ≥1.10, or MAF≥0.1% and OR≥1.60. After stringent harmonised quality control, 31 of the 32 GWAS were imputed using the 64,976 whole-genome sequenced haplotypes of the Haplotype Reference Consortium (HRC)^4^: the exception was the deCODE GWAS, which was imputed using a population-specific reference panel based on 30,440 Icelandic whole-genome sequenced haplotypes^5^ (**Methods, Supplementary Table 1**). We conducted T2D association analyses with and without adjustment for body-mass index (BMI).

### Discovery of novel loci for T2D susceptibility

We tested for T2D association with ˜27 million genotyped or imputed variants passing quality control filters, 21 million with MAF<5%. Our GWAS meta-analysis identified variants at 231 loci at genome-wide significance (*p*<5×10^−8^) in the BMI-unadjusted analysis and 152 in the smaller (effective sample size 157,401) BMI-adjusted analysis. The BMI-adjusted signals included 140 overlapping with the BMI-unadjusted analysis and 12 that were genome-wide significant only after adjustment. Of these 243 loci, 162 were significant at the more stringent threshold of *p*<5×10^−9^ recently advocated for whole genome sequence data^6^ (and therefore likely overly conservative for the GWAS setting) (**Methods**). Of the 243 significant loci, 135 mapped outside regions previously-implicated in T2D-risk, defined as >500kb from reported lead variants (**Methods, Figure 1, Supplementary Table 2**). The MAF of lead variants at the 135 novel loci ranged from 0.02% to 49.7%, including 9 low-frequency (0.5%≤MAF<5%) and 6 rare (MAF<0.5%) variant signals.

**Figure 1.**
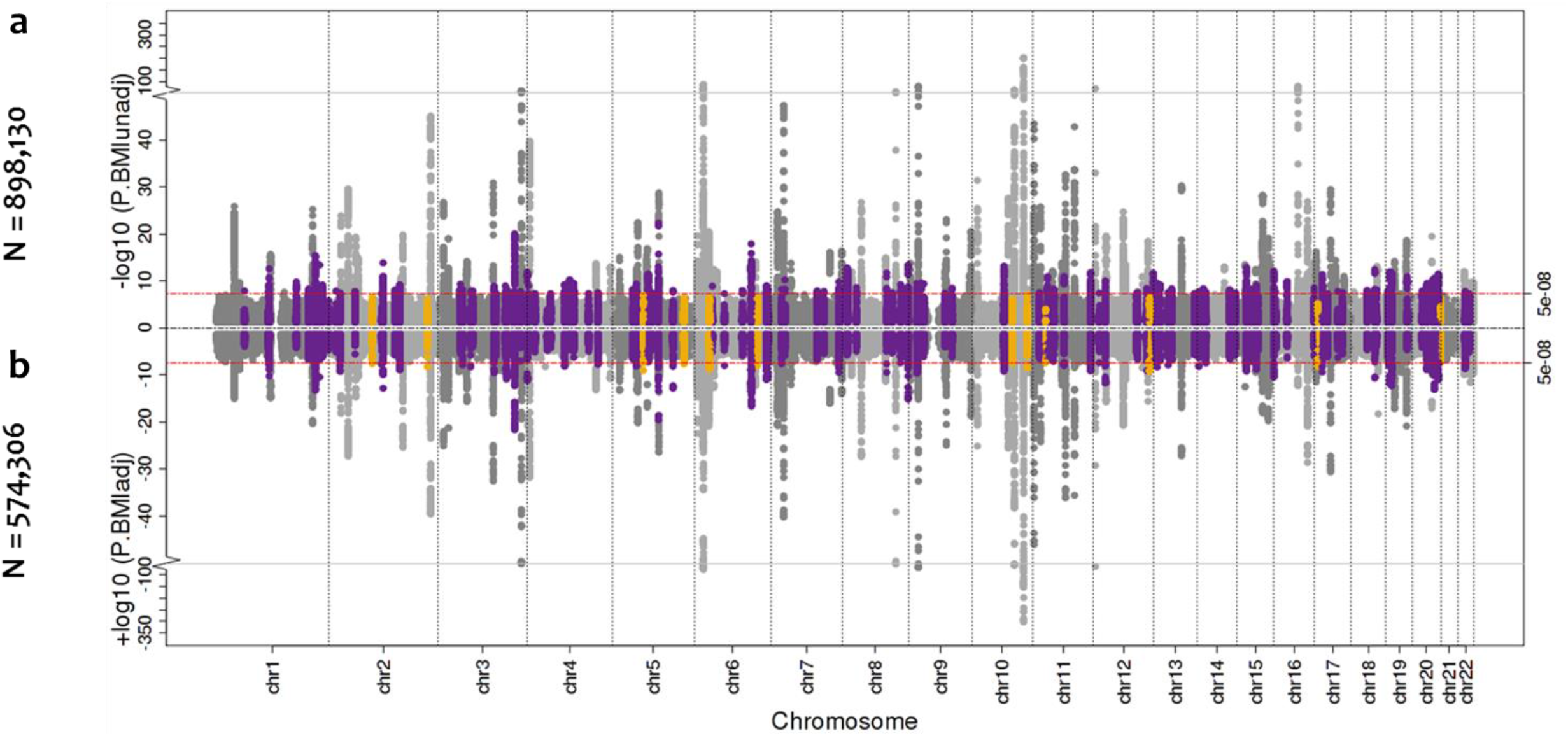
Manhattan plots of the sex-combined BMI-unadjusted and adjusted meta-analysis for T2D. **a,** Manhattan plot (top panel) of genome-wide association results for T2D without BMI adjustment from meta-analysis of up to 71,124 cases and 824,006 controls. The association *p*-value (on -log_10_ scale) for each SNP (*y*-axis) is plotted against the genomic position (NCBI Build 37; *x*-axis). Association signals that reached genome-wide significance (*p*<5×10^−8^) are shown in purple if novel. **b,** Manhattan plot (bottom panel) of genome-wide association results for T2D with BMI adjustment from meta-analysis of up to 50,409 cases and 523,897 controls. Novel association signals that reached genome-wide significance (*p*<5×10^−8^) only in the BMI-unadjusted analysis are shown in orange.

Amongst the samples not included in previous discovery efforts (42,734 cases and 497,261 controls), we replicated associations (directionally consistent at *p*<0.05) at 126 of 141 previously-reported T2D loci, including all 105 regions first discovered in European-only or trans-ethnic efforts^3,7-9^. We also detected significant European signals (*p*<0.05) for 20 other loci initially reported in studies focused on non-European subjects^10,11^. The 15 signals that did not replicate were all first identified in non-European ancestry samples: at six of these, the reported lead variant had MAF<1% in Europeans.

### Multiple association signals at T2D susceptibility loci

We used approximate conditional analysis with GCTA^12^ to delineate distinct association signals arising from multiple causal variants in the same locus (**Methods**). Across the 243 associated loci, we identified 160 additional signals at “locus-wide” significance (which, as in previous T2D analyses, we define using a locus-specific threshold of *p*<10^−5^) (**Methods**), 110 within previously-reported T2D loci. Of these 160 signals, 92 met a more conservative association threshold of *p*<10^−6^. Overall, we observed one signal at 151 loci, and two to ten signals at the remaining 92 (**Supplementary Table 2**), **for** a total of 403 significant T2D-association signals.

These analyses revealed marked complexity at some of the associated loci. For the first time, we observed evidence of multiple association signals at the *TCF7L2* locus, the largest-effect common variant signal for T2D in Europeans. In all, we detected seven additional distinct signals (0.5%<MAF<47.6%, 1.05<OR<1.36), represented by index variants that are, at most, in weak LD (r^2^<0.06) with the previously-reported lead GWAS SNP, rs7903146 (OR 1.37, MAF 29.3%). Most spectacularly, in the ˜1Mb region that encompasses the (previously-annotated) *INS-IGF2* and *KCNQ1* loci, we found compelling evidence, on conditional analyses, for 5 and 10 independent (r^2^<0.25) non-coding index variants, respectively (0.15%<MAF<42.8%,1.03<OR<1.68). The adjacency of these two loci, in a region notable for multiple imprinting effects that include many strong biological candidates (including *INS, IGF2, KCNQ1*, and *CDKN1C*), speaks to a previously-unheralded degree of complexity in this T2D-associated telomeric region of chromosome 11.

### The effects of BMI and sex

At most known and novel T2D-loci, there were only minimal differences in statistical significance and estimated T2D effect size between BMI-adjusted and unadjusted models (**Figure 2**). However, at the index SNPs for 41 distinct signals (mapping to 21 known and 16 novel loci), we observed significant heterogeneity in effect sizes between BMI adjusted and unadjusted analyses (*p*_het_<0.05/403=0.00012 adjusting for 403 variants tested; **Methods**). The heterogeneity in effect sizes followed two distinct patterns. At 26 signals, including variants at the *FTO, MC4R, TMEM18, SEC16B*, and *GNPDA2* loci, BMI-adjustment led to marked attenuation of signals detected in BMI-unadjusted analysis. These signals display strong positive correlation between BMI and T2D effect sizes, and represent T2D-risk effects primarily driven by adiposity **(Supplementary Table 3, Figure 2)**. The other 15 were more strongly associated in the BMI-adjusted analysis (including two of the 12 signals genome-wide significant in BMI-adjusted analyses alone). These reflect a mixture of signals, including some with a particularly marked effect on insulin secretion (e.g. *TCF7L2, ARAP1, JAZF1*), and others that likely influence T2D risk through reduced capacity for fat storage in peripheral adipose tissue^13^ (e.g. *GRB14, PPARG, HMGA1, ZNF664*).

**Figure 2.**
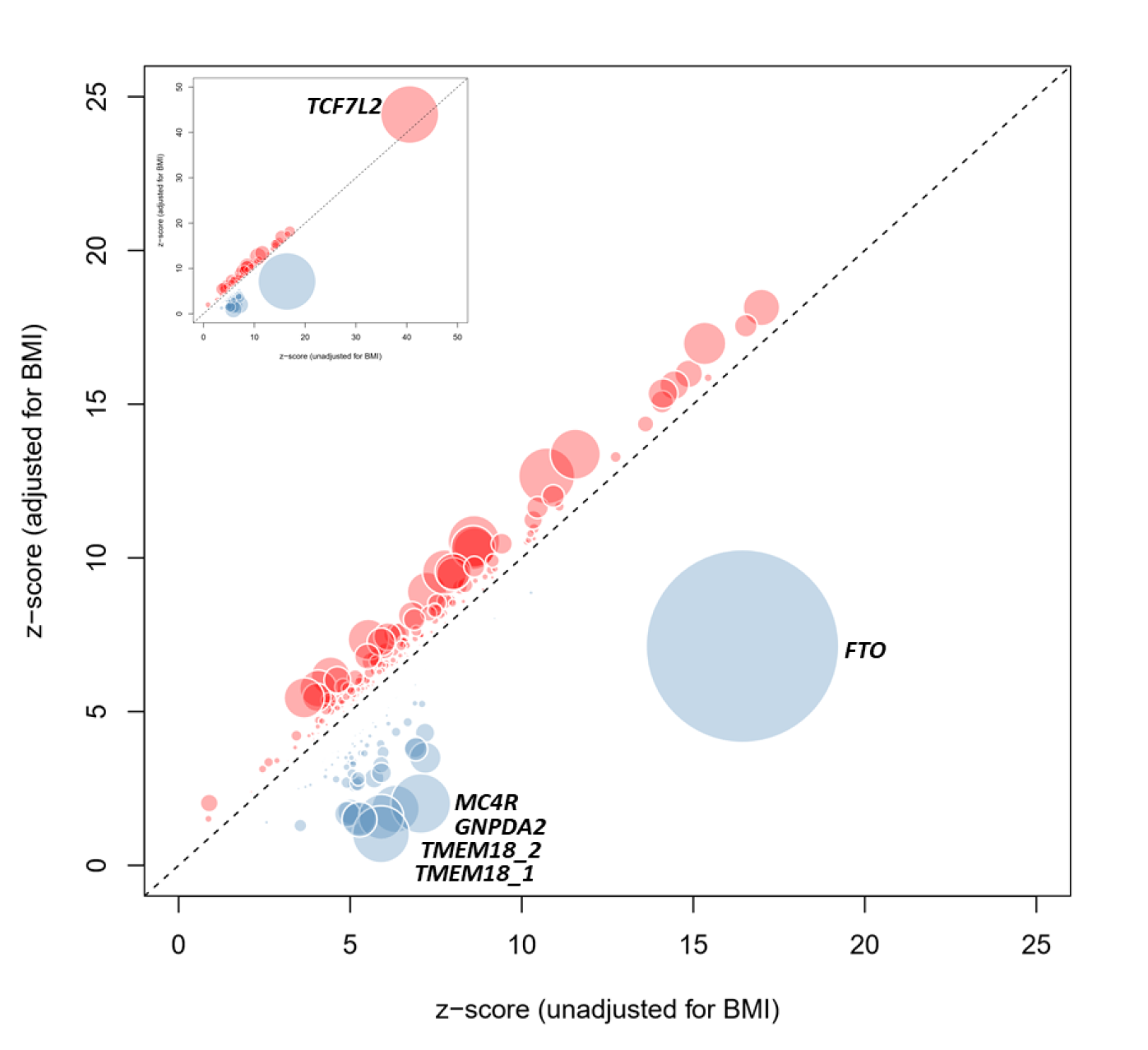
Comparison of estimated T2D effect size between BMI-adjusted and unadjusted models. Z-score for each of the 403 distinct signals from BMI-unadjusted analysis (50,791 cases and 526,121 controls; *x*-axis) is plotted against the z-score from the BMI-adjusted analysis (50,402 cases and 523,888 controls; *y*-axis). Variants that display higher T2D effect size in BMI-adjusted analysis are shown in red and variants with higher T2D effects in BMI-unadjusted analysis are shown in blue. Diameter of the circle is proportional to -log_10_ heterogeneity *p*-value.

We took advantage of the increased sample size to examine heterogeneity in allelic effects between males (41,846 T2D cases; 383,767 controls) and females (30,053 T2D cases; 434,336 controls). In sex-differentiated meta-analysis^14^ (**Methods**), we detected nominal evidence of heterogeneity (*p*<0.05) at 31 of 403 T2D susceptibility signals (7.7%, *p*_binomial_=0.013; **Supplementary Figures 1, Supplementary Table 3**). Variants at 19 signals displayed larger effects in females with the most differentiated signals at *CMIP* (rs2925979, female OR=1.09, male OR=1.03, *p_het_*=8.3×10^−6^), *KLF14* (rs1562396, female OR=1.09, male OR=1.04, *p_het_*=0.00048), and *MAP3K11* (rs1783541, female OR=1.10, male OR=1.04], *p_het_*=0.00058). The female-specific effects on T2D risk at *KLF14^15^* are consistent with those previously-reported for lipids^16^ and, given evidence that the risk variant acts as a regulator of *KLF14* expression in adipose tissue, indicate sexually-dimorphic effects on fat deposition. At the other 12 signals, larger effects were seen in males, with the differentiation most marked at *ANK1* (rs4736819, female OR=1.03, male OR=1.09, *p_het_*=0.00021). Amongst sex-differentiated signals, only that at *CMIP* survived Bonferroni correction for comparison across 403 T2D loci.

### Fine-mapping variants driving T2D association signals

Previous efforts at fine-mapping causal variants within T2D loci have been hampered by factors that are both biological (extensive LD) and technical (diverse genotyping scaffolds, relatively small reference panels for imputation). We sought to establish the extent to which the combination of increased sample size, enlarged reference panel, and harmonised variant quality control would enhance fine-mapping resolution. We performed comprehensive fine-mapping for 380 of the 403 independent T2D association signals, following conditional decomposition of loci with multiple signals (**Methods**). We excluded 23 signals not amenable to fine-mapping: (i) 19 where the index variant had MAF<0.25%; (ii) three where the index variant was rare and analysed in <50% of the effective sample size; and (iii) the one mapping to the major histocompatibility complex because of the extended and complex structure of LD across the region, which complicates fine-mapping (**Methods**). For each of the remaining 380 signals, we constructed credible sets that collectively account for ≥99% of the posterior probability of driving the association (PPA), based on the conditional meta-analysis summary statistics^17^(**Methods**).

As expected, fine-mapping resolution varied markedly across the 380 signals. Credible sets included a median of 42 variants (range 1-3,997) distributed across the entire allele-frequency spectrum (**Supplementary Figure 2**), and spanned a median of 116kb (range 1bp-995kb). At 18 signals, the credible set included a single variant; by definition, this variant had PPA>99%. For 51 signals, at 44 loci, including 18 novel loci, the most strongly associated variant accounted for >80% PPA (**Figure 3, Supplementary Table 5**).

**Figure 3.**
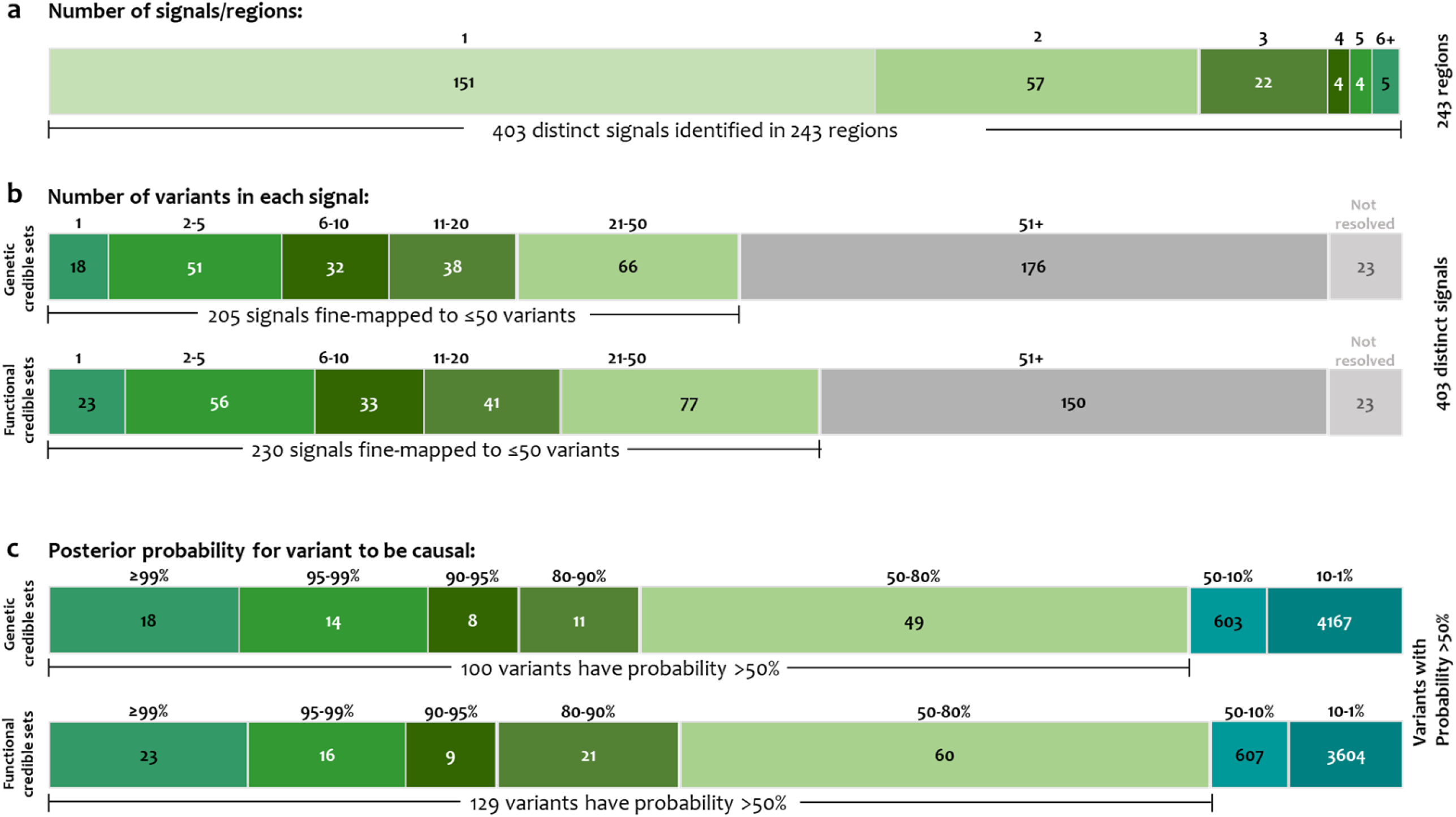
Summary of fine-mapped associations. **a,** Distinct association signals. A single signal at 151 loci, and 2–10 signals at 92. **b,** Number of variants in genetic and functional 99% credible sets. Eighteen and 23 signals were mapped to a single variant in genetic and functional credible sets, respectively. **c,** Distribution of the posterior probability of association of the variants in credible sets.

We explored fine-mapping resolution at the 83 distinct signals where detection in both studies allowed us to compare 99% credible sets from the current HRC-based analysis with those constructed using the recent meta-analysis of a subset of these T2D GWAS data imputed using the 1000G multi-ethnic reference panel^1^ (26,676 T2D cases; 132,532 controls of European ancestry, effective sample size 72,143). Genome-wide, the HRC-based meta-analysis includes 2.3-fold more variants than the 1000G study. Despite this, the HRC-imputed analysis resulted in far smaller credible sets: the median number of variants at these 83 signals decreased from 59 to 10 and the median length of intervals from 60.3kb to 19.2kb. Overall, at 79 of the 83 signals, HRC-based credible sets were either smaller (72 signals) than those generated from 1000G or unchanged (seven signals; **Figure 4, Supplementary Table 6**).

**Figure 4.**
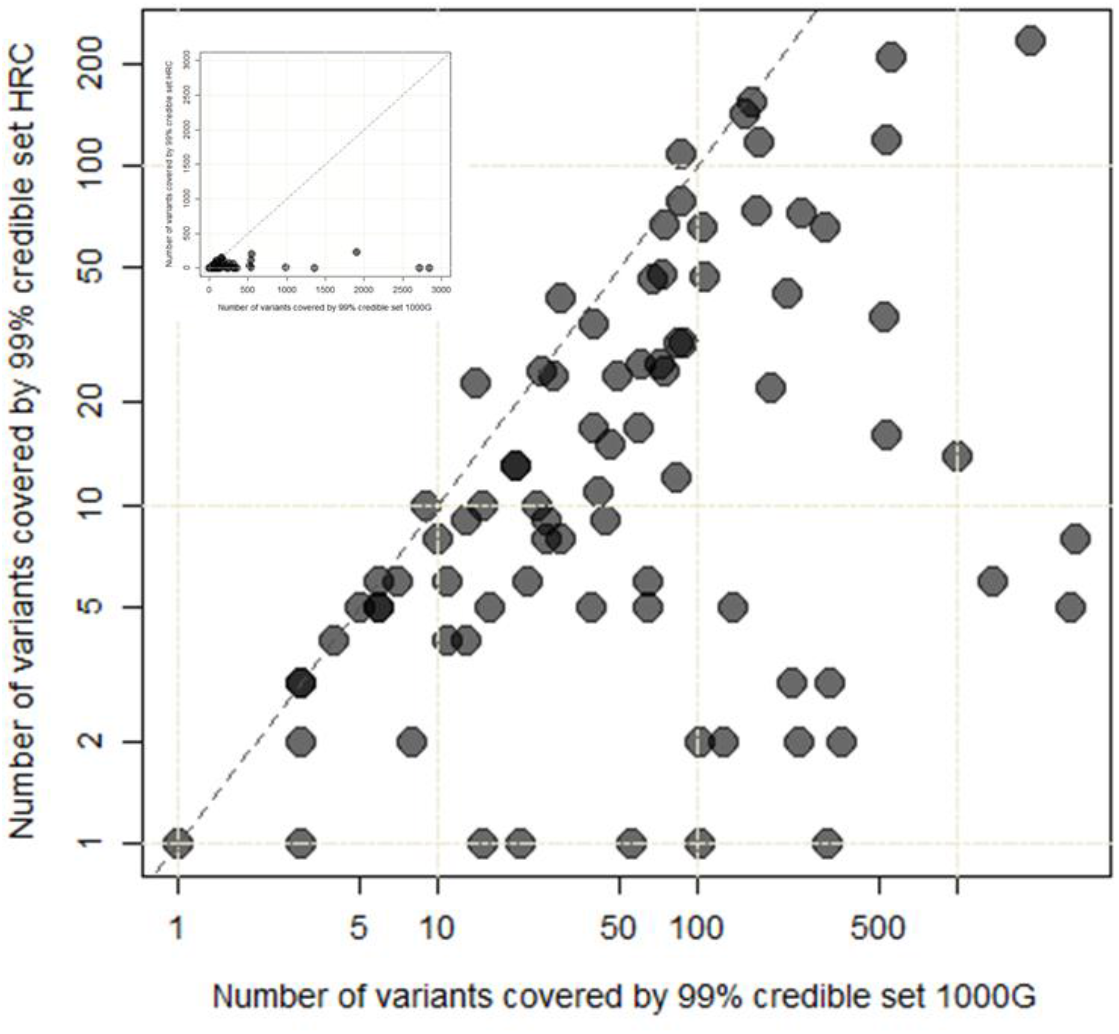
Comparison of fine-mapping resolution at 83 distinct signals. The number of variants included in the 99% credible set for each of the 83 distinct signals constructed using meta-analysis of GWAS data imputed using the 1000G multi-ethnic reference panel (26,676 T2D cases and 132,532 controls) (*x*-axis; logarithmic scale) is plotted against those (*y*-axis; logarithmic scale) derived using HRC-based imputation (74,124 T2D cases and 824,006 controls). Inset presents the same plot but with linear scales.

This improved resolution likely reflects the combination of: (i) increased effective sample size; (ii) improved imputation quality, especially for lower-frequency variants^4^; and (iii) more effective quality control measures harmonised across all contributing studies (**Methods**). To estimate the contribution to fine-mapping resolution attributable to the increase in effective sample size (the other two factors are more difficult to tease apart), we constructed 99% credible sets based on downscaling the HRC-imputation to a subset of 19 studies (31,387 cases; 326,742 controls, effective sample size 92,960) that contributed to both the 1000G and HRC-based analyses. Amongst 41 single signal loci with *p*<1×10^−5^ in this downscaled meta-analysis, estimates of credible set size (median 66) and interval (median 196kb) indicate that most of the benefits in causal variant resolution achieved in the current study derive from increased sample size.

The HRC panel provides excellent coverage of all but very rare SNVs. However, one HRC limitation is the absence of indels: these make up 4% of total variants in the phase 3 1000G reference panel^18^. To explore the potential impact on our findings, we considered the 245,207 indels from the European subset of the 1000G panel that map within 500kb of index variants at the 380 fine-mapped signals: these account for 2.8% of variants across the 380Mb of sequence. Only 1% of these are in even moderate LD (r^2^>0.5) with the index variants for each distinct signal: this suggests that indel omission is unlikely to have impacted our estimates of credible set size.

### The contribution of lower-frequency variants

The limited yield of low-frequency and rare variant signals for T2D in previous GWAS has placed an upper bound on their individual and collective contribution to disease risk^19^. The present analysis, with larger sample size and improved imputation, provided far greater power in this regard, identifying 56 low-frequency and 24 rare T2D-associated variants across 60 loci (**Figure 5**). At 21 loci, these alleles (15 low frequency, 6 rare) represented the strongest regional associations and the remaining 59 were index variants for secondary signals. Six of these 80 signals mapped within known T2D loci, five reconfirming earlier observations. At the sixth (*TMEM18*), fine-mapping highlighted the lead regional SNP as rs62107261 (MAF=4.6%) rather than the previously-reported common lead variant rs2867125 (MAF=17%)^2^; at this locus, the common and low-frequency variants constitute distinct signals, with the former relegated to secondary status in this analysis following emergence of the stronger association at rs62107261.

**Figure 5.**
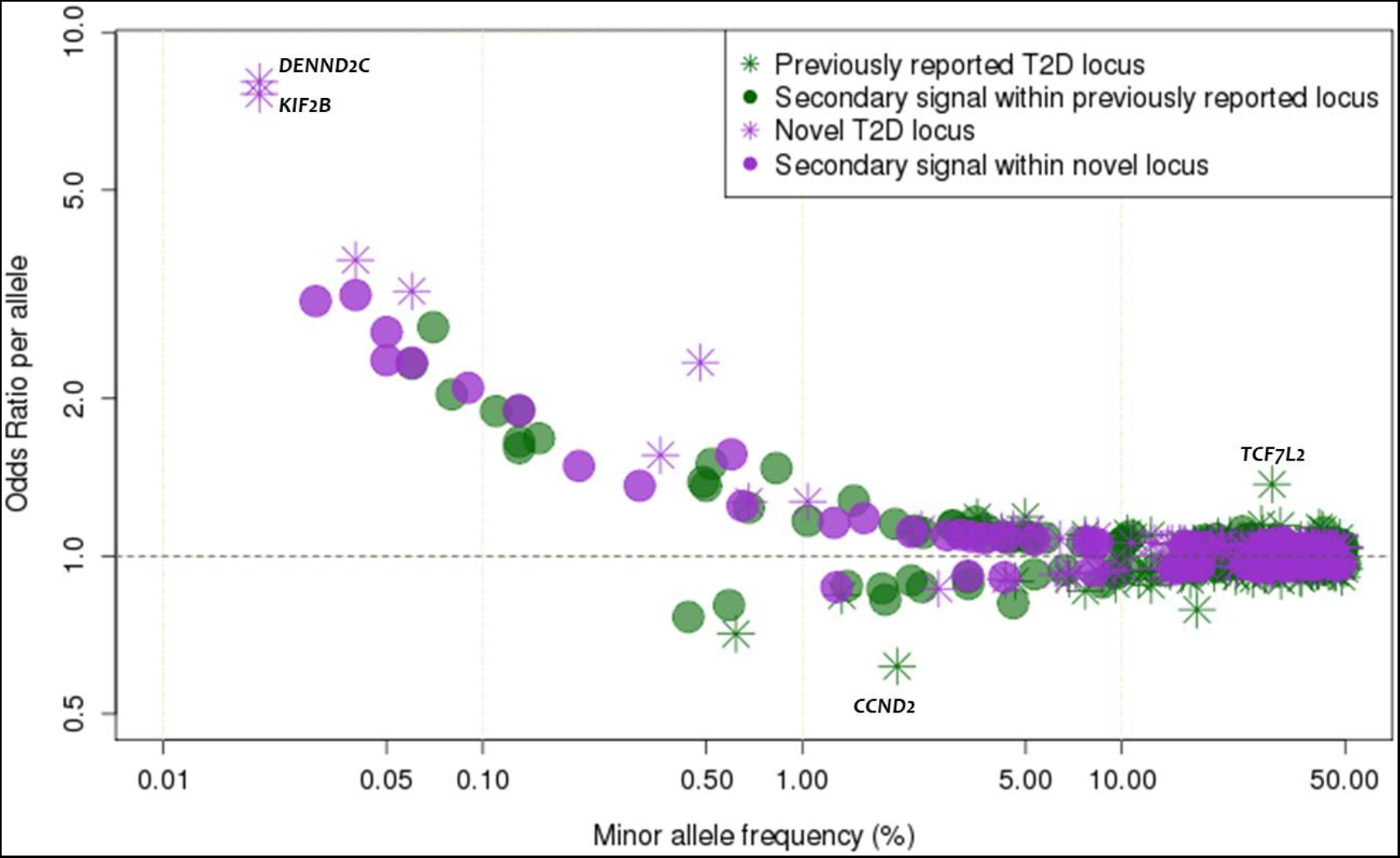
The relationship between effect size and minor allele frequency. Conditional and joint analysis effect size (*y*-axis) and minor allele frequency (*x*-axis) for 403 conditionally independent SNPs. Previously-reported T2D associated variants are shown in green and novel variants are shown in purple. Stars and circles represent the “strongest regional lead at a locus” and “lead variants for secondary signals”, respectively.

Odds ratios for low-frequency and rare variants ranged from 1.08 to 8.05 (including 14 with estimated allelic OR>2), compared to 1.03 to 1.37 for common variants (**Figure 5**). The 80 low-frequency and rare risk-variants cumulatively explain only 1.13% of phenotypic variance in T2D as compared to the 16.3% attributable to common variant signals. Extrapolation beyond these discovered signals to estimate the full contribution of lower-frequency variants to T2D risk is intrinsically difficult given the combination of effect size overestimation in the current study (winner’s curse) and limited power to capture additional low-frequency and rare variants of lesser effect. Nonetheless, these data are consistent with recently-proposed models for the genetic architecture of T2D derived from modelling findings from GWAS and sequencing data^19^. Put succinctly, although the present study leverages increased sample size and improved imputation to expand the number of low-frequency and rare variants reaching genome-wide (or for secondary signals, locus-wide) significance, the effect sizes of the associated low-frequency and rare variants are mostly modest, and this limits their collective contribution to heritability. Notwithstanding, the identification of low-frequency and rare variants of modest to large effect can provide valuable biological inference; here we briefly describe a subset of these signals.

We observed a mix of common and low-frequency variant signals around the *NEUROG3* gene, including T2D risk attributable to the minor alleles at rs41277236 (p.Gly167Arg, MAF=4.3%, OR=1.09, *p*=1.5×10^−6^) and rs549498088 (non-coding, MAF=0.60%, OR=1.56, *p*=4.7×10^−7^). *NEUROG3* encodes the neurogenin-3 transcription factor known to play a central role in the development of pancreatic islets and enteroendocrine cells^20^. In humans, rare homozygous, hypomorphic missense mutations in *NEUROG3* (non-overlapping with those we detected) are a cause of childhood-onset diabetes associated with severe congenital malabsorptive diarrhea^21^. Age of T2D diagnosis amongst heterozygous and homozygous carriers of the low-frequency *NEUROG3* T2D-risk alleles we identified was, in UK Biobank, similar to that in non-carriers (52.3 vs 52.7 years; *p*=0.21 for rs41277236; 51.1 vs 52.7 years; *p*=0.49 for rs549498088), consistent with a spectrum of functional impact that associates these variants with risk of typical T2D. To explore the phenotypic correlates of these variants, particularly with respect to the syndromic consequences of severe *NEUROG3* mutations, we conducted “PheWAS” analyses in UK Biobank (**Methods**). T2D-risk alleles at *NEUROG3* were associated with hyperlipidaemia (OR=1.07; *p*=0.0014) and multiple phenotypes recapitulating the gastrointestinal component of the neonatal syndrome (including “obstruction of bile duct” [OR=1.29; *p*=0.023], “gastrointestinal complications” [OR=1.79; *p*=0.024], and “functional digestive disorders” [OR=1.06; *p*=0.027]).

We identified a rare missense coding variant, p.Cys418Arg in *ABCC8* (rs67254669, MAF=0.11%, OR=1.89, *p*=1.1×10^−8^), which is independent of the existing GWAS lead variant in the adjacent *KCNJ11* gene (rs5213, MAF=36.2%, OR=1.07, *p*=3.5×10^−27^). *KNCJ11* and *ABCC8* encode the two components of the beta-cell K_ATP_ channel; loss and gain-of-function mutations in both genes have been reported as causal for congenital hyperinsulinemia and monogenic diabetes, respectively^22-25^. The p.Cys418Arg *ABCC8* missense variant has been previously-identified in an individual with recessively-inherited congenital hyperinsulinism^26^.

However, the relationship between hyper- and hypo-secretion of insulin associated with variation in this locus is dynamic and complex^27^: some dominant *ABCC8* mutations have been implicated in the combination of neonatal hyperinsulinism and later-onset diabetes.

We detected two previously-unreported rare alleles with large ORs. The first was intronic to *DENND2C* (rs184660829, MAF=0.020%, OR=8.05, *p*=2.5×10^−8^). In PheWAS in the UK Biobank, the T2D-risk allele was also associated with “lower gastrointestinal congenital anomalies” (OR=17.31 *p*=0.00047), and “disorders of bilirubin excretion” (OR=10.13; *p*=0.022). The second mapped near *KIF2B* (rs569511541, MAF=0.020%, OR=7.63, *p*=1.5×10^−8^) and was also associated with “congenital anomalies of endocrine gland” (OR=30.75; *p*=0.00015), “disease of pancreas” (OR=5.93; *p*=0.0017), and “hypokalemia” (OR=6.92; *p*=0.0046). Both sites are present in the Genome Aggregation Database^28^ and met quality control criteria in our data (average imputation quality >0.7; association signal visible in multiple contributing studies), but their precise contribution to T2D-risk requires further *in vitro* and *in vivo* validation.

### Causal coding variants

We next focused attention on the 51 (of 380) signals where our fine-mapping strongly implicated (PPA>80%) a single causal variant. Eight of these 51 variants were missense coding variants (**Supplementary Table 7**). Six of the eight fell into established T2D-associated regions; with the exception of p.Cys130Arg at *APOE* (MAF=15.4%), all have been previously implicated as causal variants for T2D: p.Ser539Trp in *PAM* (MAF=0.83%); p.Thr139Ile in *HNF4A* (MAF=3.5%); p.Asp1171Asn in *RREB1* (MAF=11.3%); p.Ala146Val in *HNF1A* (MAF=2.9%); and p.Pro446Leu in *GCKR* (MAF=39.3%)^3^. Coding variant associations at *PATJ* (p.Gly157Val; 9.5% MAF) and *CDKN1B* (p.Val109Gly; 23.5% MAF) are novel and highlight these genes as playing direct roles in T2D-risk. *PATJ* is highly expressed in brain^29^ and codes for Pals1-Associated Tight Junction component, a protein with multiple PDZ domains that mediate protein-protein interactions. In PheWAS, associations for this variant indicate a central mechanism of action on T2D: the T2D-risk allele is associated with obesity (OR=1.11; *p*=3.8×10^−5^; with attenuation of T2D association signal in BMI-adjusted analyses, *p*_het_=9.3×10^−10^), hyperlipidemia (OR=1.05; *p*=0.00048), and chronotype (*p*=7.7×10^−12^). *CDKN1B*, encodes a cyclin dependent kinase inhibitor; and in mouse, deletion of this gene ameliorates hyperglycemia by increasing islet mass and maintaining compensatory hyperinsulinemia^30^.

At a further four signals, a single missense variant accounted for the majority (>50%) of the PPA. These included two variants previously-reported as causal^3^: *ANKH* (p.Arg187Gln, MAF=0.6%), and *POC5* (p.His36Arg, MAF=39.5%)^3^ (**Supplementary Table 7**). The other two signals were driven by low-frequency variants: p.Gly167Arg at *NEUROG3* (described above), and p.Glu4Gln at *ZNF771* (MAF=0.6%).

### Integration of regulatory annotations to support fine-mapping

Of the 51 variants with PPA>80%, 43 mapped to regulatory sequence: 12 of these were low-frequency or rare, including variants near *ANKH, CCND2*, and *WDR72*. To characterise the regulatory impact of these variants, we overlaid them onto chromatin-state maps from T2D-relevant tissues (islets, liver, adipose, and skeletal muscle ^31-33^) and transcription factor binding sites^31,32^.

Twenty-eight of these variants mapped to islet enhancer or promoter elements; for 14, the chromatin states were islet-specific (**Supplementary Table 8, Supplementary Figure 3**). Thirteen of the 14 non-coding variants with PPA≥99% mapped to islet regulatory annotations, nine of them islet-specific. These data recapitulate previous findings implicating islet regulatory mechanisms driven by rs11257655 at the *CDC123*-*CAMKD1* locus and rs10830963 near *MTNRB1*^33-35^, and indicate similar molecular mechanisms operate at several other known T2D loci, including *IGF2BP2, ANK1, GLIS3, CDKN2B, KCNQ1, CCND2*, and *BCL2A*. Novel T2D signals near *ABCB10, FAM49A, LRFN2, CRHR2*, and *CASC11* also overlapped with islet-specific enhancer and/or promoter regions. High-PPA (i.e. PPA>80%) variants at 13, 10, and 7 signals overlapped with enhancers and/or promoters in adipose, skeletal muscle, and liver, respectively. All but four of these were also enhancers and/or promoters in islets: one signal (near *GLI2*) mapped to an adipose-specific enhancer, another (near *WDR72*) to a liver-specific enhancer, and two (near *PTGFRN* and *TSC22D2*) were located in enhancers in both adipose and skeletal muscle.

We next looked beyond loci where genetic data alone were sufficient to resolve the identity of the likely causal variant, and sought to establish whether the integration of genome-wide regulatory annotation data could refine mapping resolution where genetic localisation was less precise^33^. For these analyses, we focused on regulatory annotations from human islets because: (a) most established T2D-risk variants are considered, given patterns of association to continuous metabolic traits, to act through primary effects on beta-cell function^3,36,37^; (b) previous studies have shown that the strongest signal for regulatory enrichment at T2D association signals is seen for islet-specific regulatory elements^31,34^, a view supported by the annotation overlaps of the high-PPA (i.e. PPA>80%) variants described above; and (c) we had access to particularly extensive, high-resolution epigenomic and chromatin state annotation maps for human islets combining available histone modification and transcription factor ChIP-seq, ATAC-seq and whole genome bisulphite sequencing^33^.

Using the hierarchical modelling approach fGWAS^38^, genome-wide, we observed strong and significant (1.9 to 8.2 fold) enrichment of T2D-associated variation with respect to multiple islet enhancer and promoter states, as well as for coding sequence (with concomitant depletion of heterochromatin states) (**Methods, Supplementary Figure 4**). The joint annotation model retained strong and weak islet enhancers, strong and weak promoters, translational elongation, heterochromatin, (residual) low methylation, and coding sequence (**Methods, Supplementary Figure 4**). We used the parameters from this model as priors to redefine 99% credible sets for 380 independent T2D association signals amenable to fine-mapping (see above). We circumvented the default assumption in fGWAS of a single casual variant per locus by conducting these analyses on conditionally-decomposed data (noting that this does still allow for the possibility that the association at each conditional signal is distributed across multiple variants on a risk haplotype; **Methods**).

As expected, this integrated fine-mapping analysis boosted the PPA for variants overlapping enriched annotations (**Figure 5**, **Figure 6**). The median 99% credible set size declined from 42 to 32, credible intervals from 116kb to 100kb, and the maximum variant PPA per signal climbed by a median of 21%. The number of signals at which the lead variant PPA exceeded 80% increased from 51 to 73, with dramatic increases at some: e.g. at *GNG4* the PPA for rs291367 climbed from 24.0% to 84.2% (**Figure 3**).

**Figure 6.**
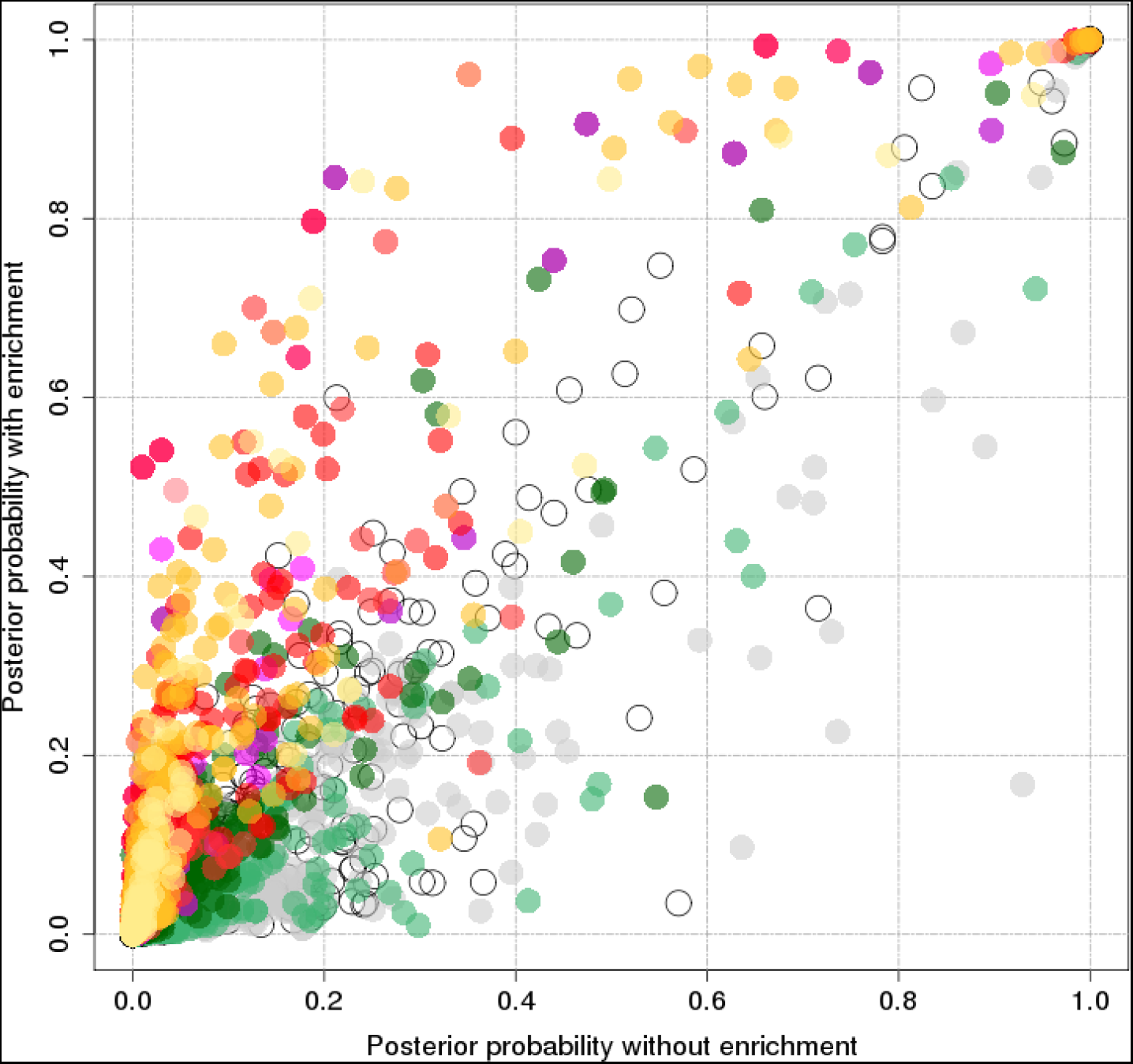
Comparison of posterior probability of association for each variant with and without incorporation enrichment information. Posterior probability of association from genetic credible sets (*y*-axis) and fGWAS analysis (*x*-axis) for each variant included in the 99% credible sets.

These annotation-supported analyses highlighted seven additional loci (further to the twelve identified on genetic evidence alone) where the majority of the PPA was invested in a coding variant (**Supplementary Table 7**). Three of these had been previously implicated as causal^3^: p.Arg276Trp at *SLC30A8* (MAF=31.5%); p.Gly226Ala at *HNF1A* (MAF=31.1%); and p.Thr113Ile at *WSCD2* (MAF=26.2%). The other four novel coding variant signals were those at *QSER1* (p.Arg1101Cys; MAF=4.3%), *SCD5* (p.Glu197Gln, MAF=33.8%), *IRS2* (p.Gly1057Asp, MAF=34.0%) and *MRPS30* (p.Glu128Gln=MAF 2.8%). In our recent study of exome array genotypes, we demonstrated that, for approximately one-third of loci harbouring coding variant associations, a causal role could be excluded once information on local LD and annotation enrichment was incorporated^3^. For all 19 coding variants (at 18 loci) describe in this study, equivalent analyses were consistent with a causal role. These analyses therefore provide multiple additional examples of human validated targets at which therapeutic modulation has the potential to modify the T2D phenotype.

Next, we concentrated on non-coding variant signals. In the fGWAS analysis, we identified 15 additional signals (beyond the 43 non-coding signals described above) at which the lead variant PPA exceeded 80% (**Supplementary Table 8**). These signals overlap active regulatory sites in islets including strong enhancers (e.g. at *TCF7L2, HNF4A, ANKH, RNF6, ZBED3*), active promoters (*EYA2*), weak enhancers (*ADSCL2, ADCY5, CDKN2B, TBCE*), and weak promoters (*DGKB*). For most of these loci, orthogonal data (e.g. associations with continuous metabolic traits^3,36,37^, *cis*-eQTL data^39^) are consistent with a role in islet function. In contrast, at six signals, including three which are likely, on physiological grounds, to be acting, at least partly, through effects on islets, we saw reductions (10% to 76%) in the lead variant PPA after islet-annotation-informed fGWAS (**Supplementary Table 8**). This arose when lead variants from the genetic fine-mapping overlapped annotations depleted in the genome-wide model. Examples included variants at the primary *CDKAL1* and secondary *KCNQ1* and *INS-IGF2* signals, where the index-variant PPA decreased by 76%, 34%, and 22%, respectively. One explanation is that these reflect T2D-association signals where the phenotypic impact on insulin secretion is mediated through the long-term consequences of regulatory influences during islet development that are no longer reflected in regulatory annotations seen in mature islets.

At many of these fine-mapped regulatory loci, the integrated data provide novel insights into disease mechanism; here we highlight three. In the T2D-association locus near *ST6GAL1*, rs3887925 achieves PPA=98.5% in the genetic data alone (99.3% in fGWAS analysis), and overlaps with enhancers active in islet, as well as liver, adipose, and skeletal muscle (**Supplementary Figure 5**). The T2D-risk allele at rs3887925 is associated with a significant increase in *ST6GAL1 cis-*expression specific to islets^39^, an effect consistent with evidence for reduced insulin secretion in risk-allele carriers during provocative testing^40^. The candidate effector transcript, *ST6GAL1*, encodes β-galactoside α2,6-sialyltransferase-1, a key enzyme responsible for the biosynthesis of α2,6-linked sialic acid in N-linked glycans. Altered glycation has the potential to impact on multiple processes, and global perturbation of *ST6GAL1* has broad effects including, in *St6gal1* knockout mice, increased body weight and visceral fat accumulation^41^. However, no equivalent association between rs3887925 and anthropometric and lipid phenotypes is seen in massive human GWAS studies^16,42,43^. This is consistent with T2D predisposition attributable to the GWAS-detected variant being mediated through regulatory mechanisms that are more tissue-specific and focused on the modulation of *ST6GAL1* expression in islets.

At the *ANK1* locus, we observed three distinct (r^2^<0.08) association signals. The strongest causal variant attribution was for the primary signal at rs13262861 (PPA=97.3% on genetic data alone; 98.8% with fGWAS). This variant overlaps an islet promoter that lies 3’ to *ANK1* but 5’ to the transcription factor *NKX6.3* (**Supplementary Figure 6**). The T2D-risk allele at the rs13262861 signal shows a directionally-consistent association with both *in vivo* measures of reduced insulin secretion^3,37,40^, and a *cis*-eQTL for reduced *NKX6-3* expression in human islets^32^. Members of the *NKX6* transcription factor family (including NKX6.3) are implicated in islet development and function^44^. A recent study describing the integration of skeletal muscle regulatory annotations with T2D association data highlighted the relationship between variants including rs515071 and rs508419 and the expression and splicing of *ANK1* in skeletal muscle^45^; however, all variants influencing *ANK1* have minimal impact on T2D-risk based on the genetic fine-mapping (PPA<1% in all three conditionally-decomposed signals). Collectively, these data indicate that the mechanism of T2D predisposition at this locus is more likely to be mediated through reduced islet expression of *NKX6.3* than altered muscle expression of *ANK1*.

At the *TCF7L2* locus, the patterns of overlap with islet and other annotations across the eight distinct T2D-association signals identified on conditional analyses offer a potential explanation for the diverse metabolic consequences of *TCF7L2* perturbation in humans and animal models (**Supplementary Table 9**)^46^. The primary signal at rs7903146, long established as the largest common variant effect for T2D in Europeans, overlaps an islet enhancer (leading to a boost in PPA from 59% to 97% on fGWAS) as well as multiple islet-relevant transcription factor binding sites, and is located in islet open chromatin^47^, all features consistent with the marked islet phenotype (deficient insulin secretion) evident for this variant in non-diabetic individuals^8^ (**Supplementary Figure 7**). However, amongst the seven secondary signals, the picture is more mixed. Amongst the four secondary signals mapped to credible sets of fewer than 10 variants, only one variant (rs144155527) rises to moderate PPA (68%) following islet annotation-enriched fGWAS analysis; many other credible set variants map to adipose and liver enhancers, illustrating the potential for their T2D-risk effects to be mediated via modulation of *TCF7L2* expression in tissues more relevant to insulin action.

### Heritability estimates and polygenic risk score prediction

Using LD score regression^48^, and empirical estimates of population- and sample-level T2D prevalence^48^, we estimated chip heritability (on the liability scale) for T2D at 18% (**Supplementary Figure 8**), a figure about half that of median estimates of heritability derived from twin and family studies^49^.

Estimates of heritability were slightly greater in females (23%) than males (17%), matching observations from a recent study of Europeans individuals based on UK Biobank alone^50^.

Identification of individuals at increased genetic risk for T2D could enhance screening strategies and allow targeted prevention. Previous attempts at using genetic data for disease prediction have shown limited utility^51,52^. We revisited this question in the present data set, using a BMI-unadjusted meta-analysis of all samples other than UK Biobank to generate genome-wide polygenic risk scores (PRS). We then applied these PRS to predict T2D status in the 18,197 cases and 423,697 controls from UK Biobank. We tested a series of PRSs, all built from 4.6M common variants but using different r^2^ and *p*-value thresholds (**Methods**)^53^ for classification performance in UK Biobank. Maximal discrimination (an AUC C-statistic of 66%: this AUC is equivalent to that obtained from BMI, age, and sex, in UK Biobank individuals) was obtained from a PRS of 136,795 variants (r^2^>0.6; *p*<0.076) (**Supplementary Figure 9**). In European subjects from UK Biobank, individuals in the top 2.5% of the PRS distribution were at 3.4-fold increased risk (prevalence=11.2%) compared to the median (prevalence=3.3%), and 9.4-fold compared to the bottom 2.5% (prevalence=1.2%). The modest T2D prevalence rates in UK Biobank reflect the age-distribution of the cohort (compared to the age at T2D onset) and preferential ascertainment of healthy individuals. If equivalent risk prediction performance applied to the general UK population, this would equate to lifetime risks for T2D of ˜51% and ˜5.5% for individuals from those extremes, based on current prevalence rates for those over 55 years of age^54^.

### Defining relationships with other traits

To characterise genetic relationships with other biomedical-relevant traits, we used LD score regression^48^ as implemented in LDHub^55^, testing 182 unique phenotypes, after excluding those with low heritability estimates and repeated measures. Eighty-five of those traits demonstrated a significant (Bonferroni corrected threshold p<0.00028) genetic correlation with T2D. A genetic increase in T2D-risk was, as expected, positively associated with multiple cardiometabolic traits including measures of adiposity, hypertriglyceridemia, coronary artery disease, and continuous glycemic traits (fasting glucose, insulin; HbA1c). (**Supplementary Table 10, Supplementary Figure 10**) and negatively correlated with HDL-cholesterol and birth weight.

These analyses also highlighted a series of more novel examples of significant genetic correlation, linking increased T2D-risk to sleeping behaviours (insomnia, excessive daytime sleeping), smoking (cigarettes smoked per day, ever versus never smoked), metabolites (glycoprotein acetyls, isoleucine, valine), depressive symptoms, urinary albumin-to-creatinine ratio, and urate. T2D-risk was negatively correlated with anorexia nervosa, intelligence, parent’s age at death, lung function measures, education status/duration, age at menarche, and age of first birth. We used the BMI-adjusted T2D-association data to demonstrate that many of these relationships (including those related to intelligence, smoking behaviour, age at menarche, age at first birth and education status) were primarily mediated by the shared impact of BMI/obesity on both T2D and the correlated phenotype **(Supplementary Figure 11)**.

## DISCUSSION

This study demonstrates how substantial increases in sample size combined with more accurate and comprehensive imputation expand characterization of the genetic contribution to T2D risk. The number of significantly associated genomic regions has doubled to ˜400, with a growing harvest of risk-alleles of lower frequency, some with relatively large effects (14 with OR>2). At many of these signals, fine-mapping resolution has been substantially improved; we mapped 51 of 380 signals to single-variant resolution on genetic evidence alone, and demonstrated that the integration of genomic annotations (here focused on those from human islets) provides further specification of plausible causal variants. We highlight 18 genes as human validated targets based on causal coding variant effects and provide novel insights into the biological mechanisms operating at several fine-mapped regulatory signals. These findings represent mechanistic hypotheses that can now be targeted for large-scale empirical validation at both the level of the variants (e.g. through massively parallel reporter assays) and the candidate effector genes (through CRISPR screens in appropriate cellular models, and manipulation in *in vivo* models). The present study was limited to individuals of European ancestry: integration of these data with large-scale GWAS data from other major ancestral groups (as is being pursued by the DIAMANTE consortium) is expected to provide an additional boost to locus discovery, and to support further increases in causal variant resolution, most obviously at loci where extensive LD within European subjects limits the power of fine-mapping.

## DATA AVAILABILITY STATEMENT

Summary level data from the exome array component of this project will be made available at the DIAGRAM consortium website http://diagram-consortium.org/ and Accelerating Medicines Partnership T2D portal http://www.type2diabetesgenetics.org/.

## ACKNOWLEDGMENTS

A full list of acknowledgments appears in the **Supplementary Information**. Part of this work was conducted using the UK Biobank resource.

## AUTHOR CONTRIBUTIONS

**Project co-ordination**. A.Mahajan, A.P.M., M.B., M.I.M. **Writing**. A.Mahajan, D.T., A.P.M., M.B., M.I.M. **Core analyses**. A.Mahajan, D.T., M.T., J.M.T., A.P.M., M.B., M.I.M. **Statistical analysis in individual studies**. A.Mahajan, D.T., N.R.R., N.W.R., V.S., R.A.S., N.G., J.P.C., E.M.S., M.W., C.Sarnowski, J.N., S.T., C.Lecoeur, M.P., B.P.P., X.G., L.F.B., J.B.-J., M.C., K.L., C.-T.L., A.E.L., J’a.L., C.Schurmann, L.Y., G.T., A.P.M. **Genotyping and phenotyping**. A.Mahajan, R.A.S., R.M., C.G., S.T., K.-U.E., K.F., S.L.R.K., F.K., I.N., C.M.B., C.Schurmann, E.P.B., I.B., C.C., G.D., I.f., V.G., M.I., M.E.J., S.L., A.L., V.L., V.M., A.D.M., G.N., N.S., A.S., D.R.W., S.S., E.P.B., S.H., C.H., J.Kriebel, T.M., A.P., B.T., D.A., G.A., C.Langenberg, N.J.W., A.P.M., M.B., M.I.M. **Islet annotations**. M.T., J.M.T., A.J.B., V.N., A.L.G., M.I.M. **Individual study design and principal investigators**. E.P.B., J.C.F., O.H.F., T.M.F., A.T.H., M.A.I., T.J., J.Kuusisto, C.M.L., K.L.M., J.S.P., K.Strauch, K.D.T., U.T., J.T., J.D., P.A.P., E.Z., R.J.F.L., P.F., E.I., L.L., L.G., M.L., F.S.C., J.W.J., C.N.A.P., H.G., A.Metspalu, D.A., A.K., G.A., J.B.M., J.I.R., J.M., O.P., T.H., C.Langenberg, N.J.W., K.Stefansson, A.P.M., M.B., M.I.M.

## MATERIALS & CORRESPONDENCE

Correspondence and requests for materials should be addressed to.

## ONLINE METHODS

### Ethics statement

All human research was approved by the relevant institutional review boards, and conducted according to the Declaration of Helsinki. All participants provided written informed consent.

### Study-level analyses

We considered a total of 74,124 T2D cases and 824,006 controls from 32 GWAS undertaken in individuals of European ancestry (**Supplementary Table 1**), genotyped with a variety of genome-wide SNP arrays. Sample and variant quality control was performed within each study (**Supplementary Table 1**). To improve the quality of the genotype scaffold in each study, we developed a harmonised protocol in which variants were subsequently removed if: (i) allele frequencies differed from those for European ancestry haplotypes from the HRC reference panel^4^ by more than 20%; AT/GC variants had MAF>40% because of potential undetected errors in strand alignment; or (iii) MAF<1% because of difficulties in calling rare variants. Each scaffold, with exception of the deCODE GWAS, was then imputed up to the HRC reference panel^4^. The GWAS from deCODE was imputed up to a reference panel based on 30,440 Icelandic whole-genome sequences^5^, and only variants that were present on the HRC panel were considered for downstream analyses. Within each study, all variants were tested for association with T2D in a regression framework, with and without adjustment for BMI, in sex-combined and sex-specific analyses, under an additive model in the effect of the minor allele, with additional adjustment for study-specific covariates (**Supplementary Table 1**). To account for population structure and relatedness, association analyses were either adjusted for principal components (after excluding related individuals) or implemented in a mixed model with random effects for kinship from a genetic relationship matrix. For studies analysed using linear mixed models, implemented in EMMAX^56^ or BOLT-LMM^57^ (**Supplementary Table 1**), allelic effects and standard errors were converted to the log-odds scale to correct for case-control imbalance^58^. For each analysis, in each study, variants were removed from a study if: (i) minor allele count <5 (in cases and controls combined); (ii) imputation quality r2-hat<0.3 (miniMAC) or proper-info<0.4 (IMPUTE4); or (iii) standard error of the allelic log-OR>10. Association summary statistics for each analysis within each study were then corrected for residual structure by means of genomic control inflation factor^59^, calculated after excluding variants mapping to established T2D susceptibility loci (**Supplementary Table 1**).

### Sex-combined meta-analysis

We aggregated association summary statistics from sex-combined analyses for each variant across studies, with and without adjustment for BMI, using fixed-effects meta-analysis with inverse-variance weighting of log-ORs, implemented in METAL^60^. The BMI unadjusted meta-analysis was subsequently corrected for residual inflation (to account for structure between studies) by means of genomic control (λ=1.013)^59^, calculated after excluding variants mapping to established T2D susceptibility loci. No adjustment was required for the BMI adjusted meta-analysis (λ=0.992). From the meta-analysis, variants were extracted that passed quality control in at least two studies. Heterogeneity in allelic effect sizes between studies contributing to the meta-analysis was assessed by Cochran’s *Q* statistic^61^. We defined novel loci as mapping >500kb and conditionally independent from a previously reported lead GWAS SNP.

For the present study, we maintained the conventional genome-wide significance threshold of 5×10^−8^, for compatibility with previous reports. We recognise that more comprehensive capture of lower-frequency variants in particular increases the effective number of tests, with some consequent increase in the false positive rate for signals just below this threshold. 162 of the 243 primary signals are significant at a more stringent threshold (5×10^−9^) recently advocated for whole genome sequence data^6^, and the major conclusions of the manuscript are unchanged if we select this more stringent (and given that our data lack the full coverage of WGS data, over-conservative) threshold. We make all summary level data results available so that readers can interpret the results themselves.

### Sex-differentiated meta-analysis

The meta-analyses described above were repeated for males and females separately, with correction for population structure by genomic control as necessary: (i) male-specific BMI unadjusted λ=1.029; (ii) male-specific BMI adjusted λ=1.001; (iii) female-specific BMI unadjusted λ=0.955; and (iv) female-specific BMI adjusted λ=0.932. The male-specific meta-analysis consisted of up to 41,846 cases and 383,767 controls, whilst the female-specific meta-analysis consisted of up to 30,053 cases and 434,336 controls. The sex-specific meta-analyses were then combined to conduct a sex-differentiated test of association and a test of heterogeneity in allelic effects between males and females^14^.

### Assessment of effect of BMI adjustment

We compared the genetic effect sizes (beta coefficients) estimated from models with and without BMI adjustments using a matched meta-analysis conducted on the same subset of 28 studies: 
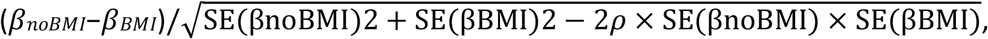
where *β_BMI_* and *β_noBMI_* are the estimated genetic effects from models with and without BMI adjustment, SE(*β*) is the estimated standard error of the estimates, and *ρ* = 0.89, is the estimated correlation between *β_BMI_* and *β_noBMI_* across all variants^1^.

### Detection of distinct association signals

We used GCTA^12^ to perform approximate conditional analyses to detect distinct association signals at each of the genome-wide significant risk loci for T2D (newly identified or confirmed, except at the major histocompatibility complex (MHC) region). GCTA performs conditional analysis using association summary statistics from GWAS meta-analysis and estimated LD from a sufficiently large reference study used in the meta-analysis. We used a reference sample of 6,000 (nearly) unrelated (pairwise relatedness <0.025) individuals of white British origin, randomly selected from the UK Biobank, to model patterns of LD between variants. The reference panel of genotypes consisted of the same 39 million variants from the HRC reference panel assessed in our GWAS, but with an additional quality control step to exclude SNPs with low imputation quality (proper-info<0.4) or deviation from Hardy-Weinberg equilibrium (*p*<1×10^−6^). For each locus, we first searched ±500kb surrounding the lead SNP (using summary statistics from BMI unadjusted or adjusted analysis, as appropriate) to ensure potential long-range genetic influences were assessed. Within a region, conditionally independent variants that reached locus-wide significance (*p*<10^−5^) were considered as index SNPs for distinct association signals. If the minimum distance between any distinct signals from two separate loci was less than 500kb, we performed additional conditional analysis taking both regions (encompassing ±500kb from both ends) and reassessed the independence of each signal.

### Fine-mapping of distinct association signals with T2D susceptibility

We considered 380 of the 403 identified distinct signals, excluding 23 that were not amenable to fine-mapping: (i) 19 signals with MAF<0.25%; (ii) three signals where the index variant was rare and analysed in <50% of the total effective sample size, defined as *N_e_* = 4 × *N_cases_* × *N_controls_*/(*N_cases_* + *N_controls_*); and (iii) the one signal in the major histocompatibility complex because of the extended and complex structure of LD across the region, which complicates fine-mapping.

For each of the remaining distinct signals, we first defined a genomic region 500kb on either side of the index variant, considering only variants with MAF>0.25% that were reported in at least 50% of the total effective sample size, thus removing those that were not well imputed in the majority of samples. We then adopted two approaches to compute 99% credible sets with 99% posterior probability of containing the causal variant: (i) using a (functionally unweighted) Bayesian approach, with the strength of evidence for association measured using the Bayes’ factor in favour of association for each variant^17,62^; and (ii) using (functionally weighted) fGWAS^38^ that reweights the association measures by using information from functional genomics data.

### (i) Genetic credible sets

For each distinct association signal, we first calculated an approximate Bayes factor^62^ in favour of association on the basis of allelic effect sizes and standard errors from the meta-analysis (using BMI-unadjusted or adjusted meta-analysis, as appropriate). For loci with a single association signal, effect sizes and standard errors from unconditional meta-analysis were used. For loci with multiple distinct association signals, these parameters were derived from the approximate conditional analysis adjusting for all other index variants in the region. Specifically, for the *j*th variant, 
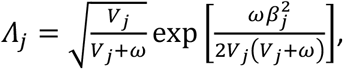
where *β_j_* and *V_j_* denote the estimated allelic effect (log-OR) and corresponding variance from the meta-analysis. The parameter *ω* denotes the prior variance in allelic effects, taken here to be 0.04^62^.

We then calculated the posterior probability that the *j*th variant drives the association signal (PPA), given by 
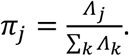

The 99% credible set^17^ for each locus was then constructed by: (i) ordering all variants in descending order of their PPA; and (ii) including ordered variants until the cumulative PPA reached 0.99. The number of variants and length of the genomic region covered by each 99% credible set was then calculated.

### (ii) Functionally weighted credible sets

We first tested each of the 15 chromatin states in human islets and coding DNA sequence separately for enrichment using genome-wide data with the program fGWAS^38^. Details on generation of the 15 chromatin states have been described elsewhere^33^. The annotation with the most significant enrichment was retained and tested jointly with each remaining annotation. If the most significant two annotation model improved the model likelihood then the two annotations in the model were retained and the process continued until the model likelihood did not exceed the previous iteration. The resulting “full” model was iteratively pruned by dropping each annotation and assessing the cross-validated likelihood of the reduced model (i.e. an annotation was removed from the “full” model if dropping it increased the cross-validated likelihood). This process resulted in the “best joint model”.

By default, fGWAS partitions the genome into “blocks” of 5,000 SNPs and assumes no more than one causal variant per block. However, for direct comparison with the “genetic” credible sets and to account for multiple distinct association signals within a locus, we used a modified approach. For T2D associated regions with no evidence of more than one distinct signal, we delineated 1Mb windows comprising all SNPs within 500 kb of the index variant and partitioned the intervening regions into ˜1Mb windows. These windows were manually input into fGWAS using the --bed command and a separate fGWAS analysis was performed using only the set of annotations remaining in the “best joint model”. The genome-wide enrichments were used as priors in a Bayesian fine-mapping analysis implemented in fGWAS to calculate posterior probabilities for each SNP in the designated windows. For the remaining regions with evidence of two or more distinct association signals, we used the results from the approximate conditional analyses described above and similarly performed a manually partitioned fGWAS analysis. We then constructed 99% credible sets as described above.

### PheWAS analyses

We performed phenome-wide association studies (PheWAS) using genotype and phenotype data from electronic health records (EHRs) from the UK Biobank. Hierarchical phenotype codes from EHRs were curated by grouping International Classification of Disease, Ninth Revision (ICD-9) clinical/billing codes as previously described^63^. Only phenotype codes with 20 or more cases and with minor allele count of 5 or greater in cases and controls were considered eligible for analysis. Logistic regression analyses were performed in individuals of European ancestry for relevant phenotype-genotype combination adjusting for six genetic ancestry principal components, array, and sex.

### Estimation of genetic variance explained

We used Linkage Disequilibrium Score Regression (LDSR)^48^ to estimate the proportion of variance explained by common genetic variants for T2D on the liability scale. As advised by the developers, these estimates were based on summary statistics (without any genomic control correction) of variants restricted to the subset of HapMap^64^ variants after excluding the MHC region. Estimations were done for both sex-combined and sex-specific (BMI-unadjusted) analyses, assuming population prevalence of 10%.

### Estimating phenotypic variance explained by SNPs

We used UK Biobank samples (19,119 T2D cases and 423,698 controls) to calculate variance explained by genome-wide significant variants. We ran a model regressing T2D status on all independently associated rare and low-frequency variants, assuming an additive model (and adjusting for sex, age, array, and 6 principal components). A separate model was run to determine the variance captured by the independently associated common variants.

### Polygenic risk score analyses

Polygenic risk score (PRSs) were created in UK Biobank samples using raw genotype data using the software PRsice^53^ using the GWAS summary statistics from the sex-combined BMI-unadjusted T2D meta-analysis excluding UK Biobank samples. PRSs were created using *p*-value thresholds ranging from 5×10^−8^ to 0.5 using LD pruning parameters of *r*^2^ 0.2 to 0.8 over 250 kb windows.

## URLs

UK Biobank, http://www.ukbiobank.ac.uk/; MACH,

http://csg.sph.umich.edu//abecasis/MaCH/;

SHAPEIT, https://mathgen.stats.ox.ac.uk/genetics_software/shapeit/shapeit.html;

GTEx, http://www.gtexportal.org/home/; Locuszoom,

http://locuszoom.sph.umich.edu/locuszoom/; 1000 Genomes Project,

http://www.1000genomes.org/; HapMap project, http://hapmap.ncbi.nlm.nih.gov/; HRC,

http://www.haplotype-reference-consortium.org/; GCTA,

http://cnsgenomics.com/software/gcta/; LDSC, https://github.com/bulik/ldsc/; LD Hub,

http://ldsc.broadinstitute.org/; bedtools, http://bedtools.readthedocs.io/en/latest/;

DIAGRAM Consortium, http://diagram-consortium.org/; fGWAS

https://github.com/joepickrell/fgwas.

## REFERENCES

1. Scott, R.A. et al. An Expanded genome-wide association study of type 2 diabetes in Europeans. Diabetes 66, 2888–2902 (2017).

2. Zhao, W. et al. Identification of new susceptibility loci for type 2 diabetes and shared etiological pathways with coronary heart disease. Nat Genet 49, 1450–1457 (2017).

3. Mahajan, A. et al. Refining the accuracy of validated target identification through coding variant fine-mapping In type 2 diabetes. Nat Genet (accepted).

4. McCarthy, S. et al. A reference panel of 64,976 haplotypes for genotype imputation. Nat Genet 48, 1279–83 (2016).

5. Jonsson, H. et al. Whole genome characterization of sequence diversity of 15,220 Icelanders. Sci Data 4, 170115 (2017).

6. Pulit, S.L., de With, S.A. & de Bakker, P.I. Resetting the bar: Statistical significance in whole-genome sequencing-based association studies of global populations. Genet Epidemiol 41, 145–151 (2017).

7. Flannick, J. & Florez, J.C. Type 2 diabetes: genetic data sharing to advance complex disease research. Nat Rev Genet 17, 535–49 (2016).

8. Voight, B.F. et al. Twelve type 2 diabetes susceptibility loci identified through large-scale association analysis. Nat Genet 42, 579–89 (2010).

9. Morris, A.P. et al. Large-scale association analysis provides insights into the genetic architecture and pathophysiology of type 2 diabetes. Nat Genet 44, 981–90 (2012).

10. Kooner, J.S. et al. Genome-wide association study in individuals of South Asian ancestry identifies six new type 2 diabetes susceptibility loci. Nat Genet 43, 984–9 (2011).

11. Cho, Y.S. et al. Meta-analysis of genome-wide association studies identifies eight new loci for type 2 diabetes in east Asians. Nat Genet 44, 67–72 (2011).

12. Yang, J. et al. Conditional and joint multiple-SNP analysis of GWAS summary statistics identifies additional variants influencing complex traits. Nat Genet 44, 369–75, s1–3 (2012).

13. Lotta, L.A. et al. Integrative genomic analysis implicates limited peripheral adipose storage capacity in the pathogenesis of human insulin resistance. Nat Genet 49, 17–26 (2017).

14. Magi, R., Lindgren, C.M. & Morris, A.P. Meta-analysis of sex-specific genome-wide association studies. Genet Epidemiol 34, 846–53 (2010).

15. Small, K.S. et al. Regulatory variants at *KLF14* influence type 2 diabetes risk via a female-specific effect on adipocyte size and body composition. Nat Genet (accepted).

16. Teslovich, T.M. et al. Biological, clinical and population relevance of 95 loci for blood lipids. Nature 466, 707–13 (2010).

17. Maller, J.B. et al. Bayesian refinement of association signals for 14 loci in 3 common diseases. Nat Genet 44, 1294–301 (2012).

18. Auton, A. et al. A global reference for human genetic variation. Nature 526, 68–74 (2015).

19. Fuchsberger, C. et al. The genetic architecture of type 2 diabetes. Nature 536, 41–7 (2016).

20. Gradwohl, G., Dierich, A., LeMeur, M. & Guillemot, F. neurogenin3 is required for the development of the four endocrine cell lineages of the pancreas. Proc Natl Acad Sci U S A 97, 1607–11 (2000).

21. Rubio-Cabezas, O. et al. Permanent Neonatal Diabetes and Enteric Anendocrinosis Associated With Biallelic Mutations in NEUROG3. Diabetes 60, 1349–53 (2011).

22. Babenko, A.P. et al. Activating mutations in the ABCC8 gene in neonatal diabetes mellitus. N Engl J Med 355, 456–66 (2006).

23. Gloyn, A.L. et al. Activating mutations in the gene encoding the ATP-sensitive potassium-channel subunit Kir6.2 and permanent neonatal diabetes. N Engl J Med 350, 1838–49 (2004).

24. Aguilar-Bryan, L. et al. Cloning of the beta cell high-affinity sulfonylurea receptor: a regulator of insulin secretion. Science 268, 423–6 (1995).

25. Thomas, P., Ye, Y. & Lightner, E. Mutation of the pancreatic islet inward rectifier Kir6.2 also leads to familial persistent hyperinsulinemic hypoglycemia of infancy. Hum Mol Genet 5, 1809–12 (1996).

26. Aguilar-Bryan, L. & Bryan, J. Molecular biology of adenosine triphosphate-sensitive potassium channels. Endocr Rev 20, 101–35 (1999).

27. Huopio, H. et al. A new subtype of autosomal dominant diabetes attributable to a mutation in the gene for sulfonylurea receptor 1. Lancet 361, 301–7 (2003).

28. Lek, M. et al. Analysis of protein-coding genetic variation in 60,706 humans. Nature 536, 285–91 (2016).

29. Human genomics. The Genotype-Tissue Expression (GTEx) pilot analysis: multitissue gene regulation in humans. Science 348, 648–60 (2015).

30. Uchida, T. et al. Deletion of Cdkn1b ameliorates hyperglycemia by maintaining compensatory hyperinsulinemia in diabetic mice. Nat Med 11, 175–82 (2005).

31. Pasquali, L. et al. Pancreatic islet enhancer clusters enriched in type 2 diabetes risk-associated variants. Nat Genet 46, 136–143 (2014).

32. Varshney, A. et al. Genetic regulatory signatures underlying islet gene expression and type 2 diabetes. Proc Natl Acad Sci U S A 114, 2301–2306 (2017).

33. Thurner, M. et al. Integration of human pancreatic islet genomic data refines regulatory mechanisms at Type 2 Diabetes susceptibility loci. bioRxiv (2017).

34. Gaulton, K.J. et al. Genetic fine mapping and genomic annotation defines causal mechanisms at type 2 diabetes susceptibility loci. Nat Genet 47, 1415–25 (2015).

35. Fogarty, M.P., Cannon, M.E., Vadlamudi, S., Gaulton, K.J. & Mohlke, K.L. Identification of a regulatory variant that binds FOXA1 and FOXA2 at the CDC123/CAMK1D type 2 diabetes GWAS locus. PLoS Genet 10, e1004633 (2014).

36. Dimas, A.S. et al. Impact of type 2 diabetes susceptibility variants on quantitative glycemic traits reveals mechanistic heterogeneity. Diabetes 63, 2158–71 (2014).

37. Wood, A.R. et al. A Genome-Wide Association Study of IVGTT-Based Measures of First-Phase Insulin Secretion Refines the Underlying Physiology of Type 2 Diabetes Variants. Diabetes 66, 2296–2309 (2017).

38. Pickrell J, K. Joint Analysis of Functional Genomic Data and Genome-wide Association Studies of 18 Human Traits. Am J Hum Genet 94, 559–73 (2014).

39. van de Bunt, M. et al. Transcript expression data from human islets links regulatory signals from genome-wide association studies for type 2 diabetes and glycemic traits to their downstream effectors. PLoS Genet 11, e1005694 (2015).

40. Prokopenko, I. et al. A central role for GRB10 in regulation of islet function in man. PLoS Genet 10, e1004235 (2014).

41. Kaburagi, T., Kizuka, Y., Kitazume, S. & Taniguchi, N. The Inhibitory Role of alpha2,6-Sialylation in Adipogenesis. J Biol Chem 292, 2278–2286 (2017).

42. Locke, A.E. et al. Genetic studies of body mass index yield new insights for obesity biology. Nature 518, 197–206 (2015).

43. Shungin, D. et al. New genetic loci link adipose and insulin biology to body fat distribution. Nature 518, 187–96 (2015).

44. Lizio, M. et al. Mapping mammalian cell-type-specific transcriptional regulatory networks using KD-CAGE and ChIP-seq data in the TC-YIK cell line. Front Genet 6, 331 (2015).

45. Scott, L.J. et al. The genetic regulatory signature of type 2 diabetes in human skeletal muscle. Nat Commun 7, 11764 (2016).

46. McCarthy, M.I., Rorsman, P. & Gloyn, A.L. TCF7L2 and diabetes: a tale of two tissues, and of two species. Cell Metab 17, 157–9 (2013).

47. Gaulton, K.J. et al. A map of open chromatin in human pancreatic islets. Nat Genet 42, 255–9 (2010).

48. Bulik-Sullivan, B. et al. An atlas of genetic correlations across human diseases and traits. Nat Genet 47, 1236–41 (2015).

49. Meigs, J.B., Cupples, L.A. & Wilson, P.W. Parental transmission of type 2 diabetes: the Framingham Offspring Study. Diabetes 49, 2201–7 (2000).

50. Ge, T., Chen, C.Y., Neale, B.M., Sabuncu, M.R. & Smoller, J.W. Phenome-wide heritability analysis of the UK Biobank. PLoS Genet 13, e1006711 (2017).

51. Meigs, J.B. et al. Genotype score in addition to common risk factors for prediction of type 2 diabetes. N Engl J Med 359, 2208–19 (2008).

52. Weedon, M.N. et al. Combining information from common type 2 diabetes risk polymorphisms improves disease prediction. PLoS Med 3, e374 (2006).

53. Euesden, J., Lewis, C.M. & O’Reilly, P.F. PRSice: Polygenic Risk Score software. Bioinformatics 31, 1466–8 (2015).

54. UK government. Adult obesity and type 2 diabetes. (https://www.gov.uk/government/uploads/system/uploads/attachment_data/file/338934/Adult_obesity_and_type_2_diabetes_.pdf).

55. Zheng, J. et al. LD Hub: a centralized database and web interface to perform LD score regression that maximizes the potential of summary level GWAS data for SNP heritability and genetic correlation analysis. Bioinformatics 33, 272–279 (2017).

## References

56. Kang, H.M. et al. Variance component model to account for sample structure in genome-wide association studies. Nat Genet 42, 348–54 (2010).

57. Loh, P.R. et al. Efficient Bayesian mixed-model analysis increases association power in large cohorts. Nat Genet 47, 284–90 (2015).

58. Cook, J.P., Mahajan, A. & Morris, A.P. Guidance for the utility of linear models in meta-analysis of genetic association studies of binary phenotypes. Eur J Hum Genet 25, 240–245 (2017).

59. Devlin, B. & Roeder, K. Genomic control for association studies. Biometrics 55, 997–1004 (1999).

60. Willer, C.J., Li, Y. & Abecasis, G.R. METAL: fast and efficient meta-analysis of genomewide association scans. Bioinformatics 26, 2190–1 (2010).

61. Ioannidis, J.P., Patsopoulos, N.A. & Evangelou, E. Heterogeneity in meta-analyses of genome-wide association investigations. PLoS One 2, e841 (2007).

62. Wakefield, J. A Bayesian measure of the probability of false discovery in genetic epidemiology studies. Am J Hum Genet 81, 208–27 (2007).

63. Denny, J.C. et al. PheWAS: demonstrating the feasibility of a phenome-wide scan to discover gene-disease associations. Bioinformatics 26, 1205–10 (2010).

64. Frazer, K.A. et al. A second generation human haplotype map of over 3.1 million SNPs. Nature 449, 851–61 (2007).

